# Mustard-DB: The Database of Cellular, Molecular, and Phenotypic Data in Sulphur Mustard Exposure

**DOI:** 10.1101/2025.03.02.639327

**Authors:** Ali Ahmadi, Fatemeh Rafiei, Mohammad Alizadeh, Ahmad Chitsaz, Simin Ghasemi, Hasan Bagheri, Ali Ghazvini, Masoud Arabfard, Sadegh Azimzadeh Jamalkandi, Mostafa Ghanei

## Abstract

The significance of sulfur mustard as a chemical hazard cannot be overstated. Its potential for causing severe morbidity and long-term health issues makes understanding its effects crucial. Mustard-DB is an essential database that consolidates extensive cellular, molecular, and clinical data on sulfur mustard exposure. It stands as a critical tool for the scientific community, shedding light on the pathophysiological mechanisms and potential therapeutic interventions for those affected by this noxious agent. The database provides a detailed account of the acute and chronic cellular responses, and clinical outcomes associated with sulfur mustard in *in vitro, ex vivo* and *in vivo* studies. By offering a multidimensional view of the data, Mustard-DB enables a comprehensive understanding of the agent’s impact on human health. This manuscript presents the development, structure, and application of Mustard-DB (www.Mustard-DB.com), emphasizing its role in enhancing our response to chemical threats and improving patient outcomes.

## 2 Introduction

Sulfur mustard, a potent vesicant and chemical warfare agent, has been a subject of concern due to its deleterious effects on human health [1, 2]. The compound, bis(2-chloroethyl) sulfide, commonly known as mustard gas, has a notorious history dating back to its use in World War I and Iraq-Iran imposed war. Despite international efforts to control and eliminate such agents, the threat of sulfur mustard exposure remains, particularly in conflict zones and through accidental releases.

The pathophysiology of sulfur mustard is complex, involving multiple organ systems and leading to a range of acute and chronic health issues. The cellular and molecular mechanisms underpinning these effects are intricate, involving DNA damage, oxidative stress, and inflammatory responses, which can lead to long-term complications such as respiratory diseases, eye complications, and skin scarring [3, 4].

Given the severity of its impact, there is a pressing need for a comprehensive understanding of sulfur mustard’s effects at various biological levels and the development of effective treatments. Mustard-DB, a database dedicated to the cellular, molecular, and phenotype data related to sulfur mustard exposure, addresses this need by providing an integrated platform for researchers and healthcare professionals..

This manuscript introduces Mustard-DB, detailing its inception, structure, and the methodologies employed to curate and standardize the wealth of data it contains. It also highlights the database’s potential to facilitate research that can lead to breakthroughs in medical treatments and preventive measures for those affected by sulfur mustard exposure. Through Mustard-DB, we aim to contribute to the global effort to mitigate the health consequences of chemical warfare agents and support the victims.

## 3 Materials and methods

Mustard-DB was meticulously designed to capture a wide array of data types, including cellular, molecular (transcripts, proteins, and metabolites), and phenotype information (Figure 1).

**Figure 1.**
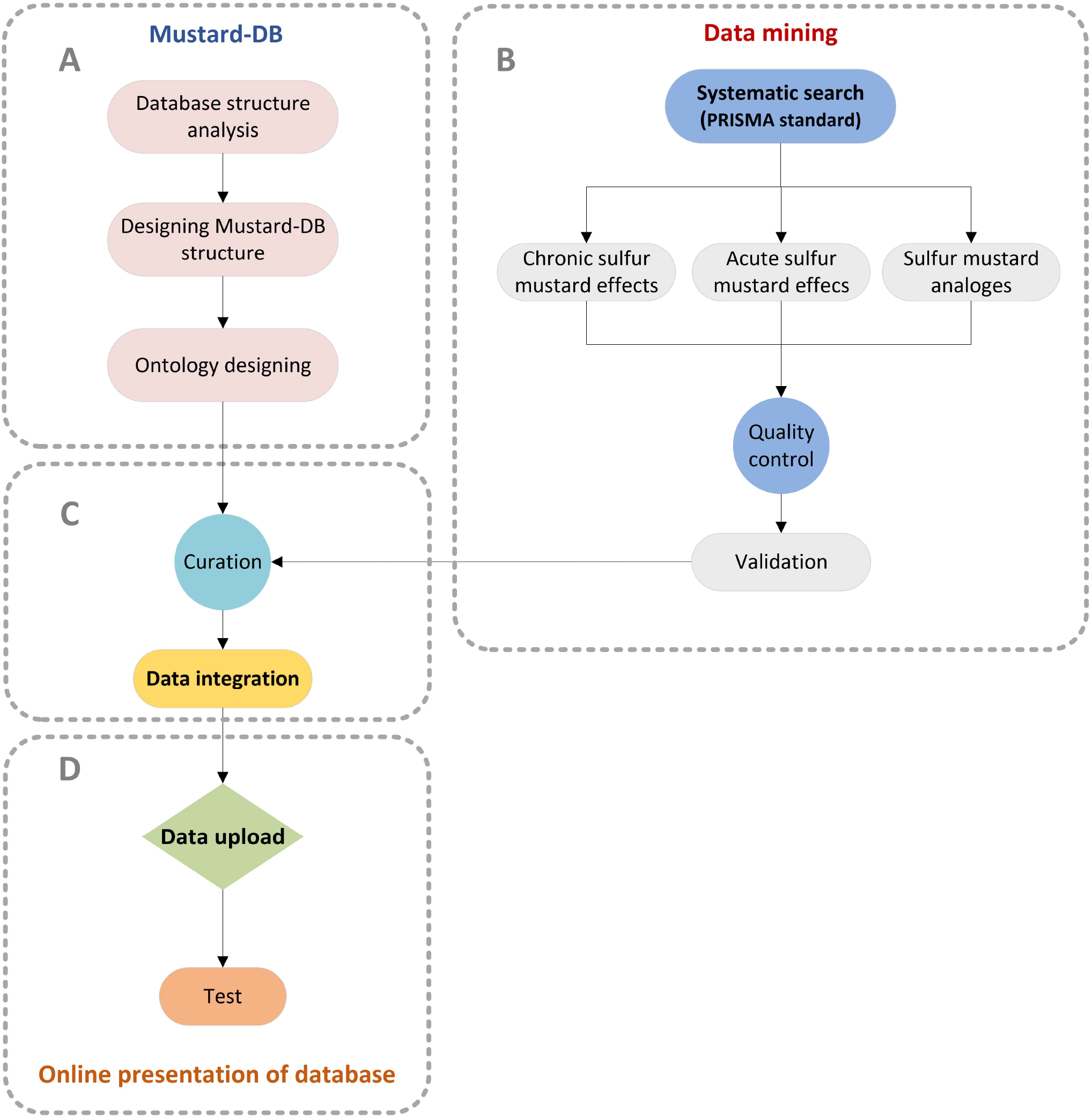
The workflow of Mustard-DB design and development includes data acquisition, database structure, and database development.

### 3.1 Database Design and Development

In the construction of Mustard-DB, a state-of-the-art technological stack was employed to ensure robustness and efficiency. The backend was developed using Laravel Version 10, renowned for its elegant syntax and comprehensive features that streamline the development process. PHP Version 8.2 was chosen for server-side scripting, seamlessly integrated within HTML to create dynamic web content. The frontend was crafted with HTML, styled with CSS for layout and design, and enhanced with JavaScript for interactivity, adhering to the principles of responsive web design.

The architecture of the database was founded on the MVC design pattern, which facilitates the separation of concerns, allowing for modular development and ease of maintenance. MySQL, a widely used relational database management system, was selected for data management due to its reliability and scalability.

## 4 Systematic search

### 4.1 Data Collection and Screening

To select eligible studies on the effects of mustard gas, the PubMed database was searched up to January 2024 using the following related terms: “Gas, Mustard” or “Di-2-chloroethyl Sulfide” or “Di 2 chloroethyl Sulfide” or “Dichlorodiethyl Sulfide” or “Sulfur Mustard” or “Yellow Cross Liquid” or “Yperite” or “Bis(beta-chloroethyl) Sulfide” or “Mustardgas” or “Psoriazin”. After the removal of duplicate articles, non-English studies, and those not available were excluded.

In the next step, review studies, studies related to yperite, and those concerning the Brassica plant were eliminated, and the remaining sources were divided into several groups based on the site of effect, namely “eyes, skin, and lungs”. Meanwhile, studies related to other body organs that were relevant to the main question were placed in separate categories. In the second screening phase, studies related to treatment, polymorphism, decontamination, soil, sensors, and diagnostics were excluded as criteria for elimination. The titles and abstracts of the remaining studies were evaluated, and ultimately, 549 studies were selected for further assessment. Figure 2 provides an overview of the systematic review of the sources. PRISMA standard 2024 was selected for further assessment [5].

**Figure 2.**
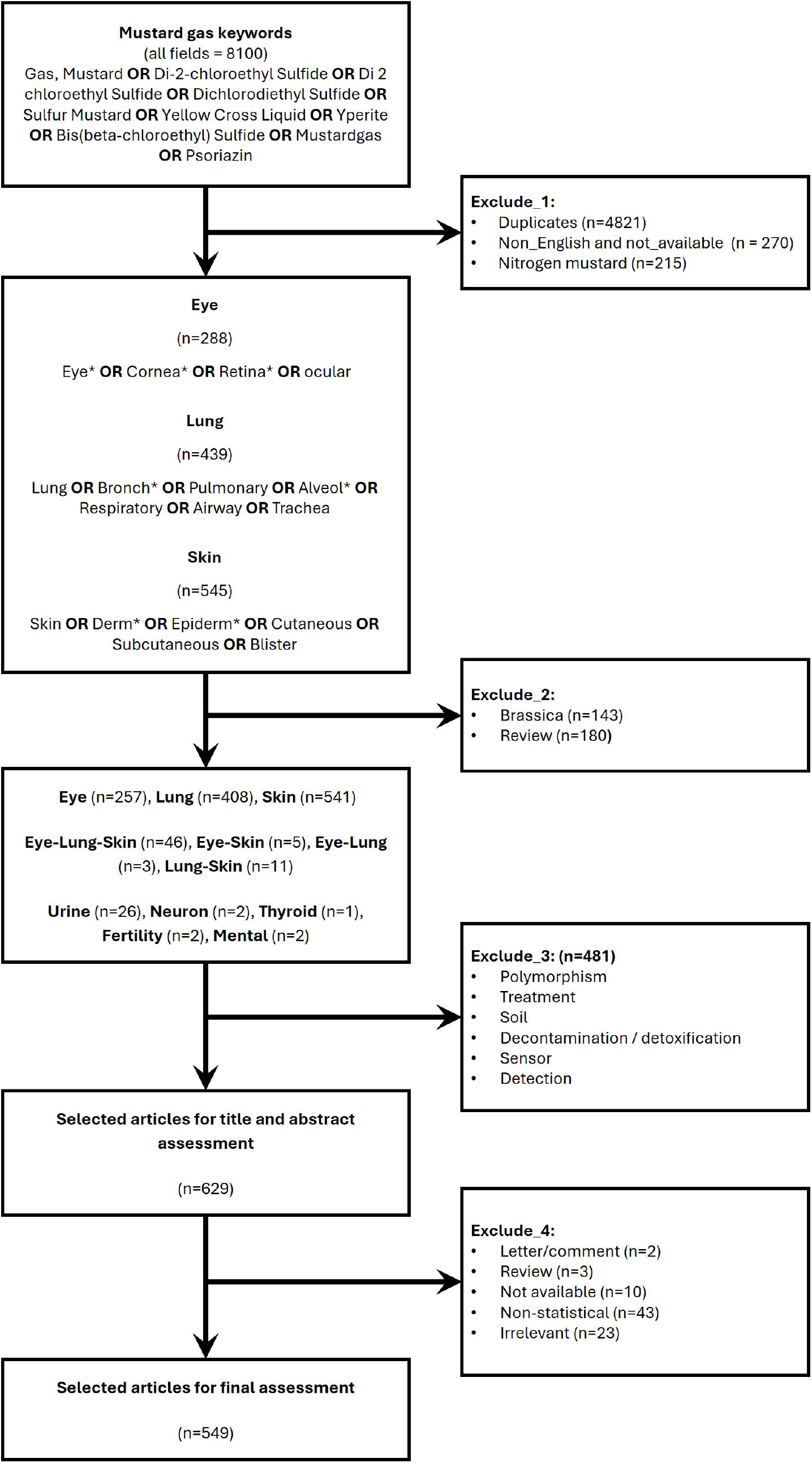
Systematic search strategy according to PRISMA quidelines.

### 4.2 Data Standardization

In the first stage, the items in question for extraction were reviewed, and 27 items for differential and 34 items for correlations/associations were selected for data extraction. In the pursuit of data standardization for Mustard-DB, a careful approach was adopted, employing a variety of ontologies to ensure uniformity and precision in data entry. The gene and protein nomenclature were aligned with the NCBI gene database, providing a comprehensive list of genes, proteins, and their alternate names. Metabolite information was standardized using the MetaboAnalyst database [6], which served as a reference for metabolite names. Cell types, sample types, and the names of organisms and diseases were categorized using established ontologies during this study to maintain consistency across the database.

Furthermore, statistical tests were codified based on the STATO ontology, a resource dedicated to the coherent representation of statistical methods [7]. The types of studies were classified according to their statistical design, ensuring clarity and ease of interpretation for database users. All references were meticulously documented, providing a trail for verification and further exploration. The adherence to these ontological standards underscores the database’s commitment to reliability and validity in the integration and presentation of sulfur mustard-related data.

### 4.3 Data entry and curation

To compile our dataset, we excluded studies that lacked statistical analysis or did not report any thresholds, such as p-values or confidence intervals. Additionally, demographic parameters were not included in the data collection as relevant variables.

Subsequently, each study was evaluated in terms of the type of condition under investigation, the type of study and experiment conducted, the type of sample, the variables of interest and their types, and matched and assessed with the ontology. Next, the type of statistical test, the p-value, and the sign of difference of the variable (increase, decrease, or no change) were entered, and the related data were extracted. Additionally, the type of laboratory technique used to measure the amount of change was considered. For data collection, Google Sheets was utilized for coding and integrating ontological information for data entry. An independent Google Sheet was designed for each user. A portion of the data is presented in Supplemental File 1. In addition to designing a dropdown menu, conditions were established to indicate whether entries were repetitive/non-repetitive and whether the data were permissible or not. Quality control of the data entry procedure was performed by two independent staff.

In compiling the results for Mustard-DB, a comprehensive approach was taken to include all relevant findings from the studies. This encompassed both significant and nonsignificant results, as well as observed changes or lack thereof in the comparisons. Such inclusivity ensures that the database provides a complete picture of the research conducted on sulfur mustard exposure, allowing users to draw their own conclusions from the full spectrum of data to support a broader range of research inquiries.

### 4.4 Data Integration

The integration of data within Mustard-DB was a critical step in providing a holistic view of the effects of sulfur mustard exposure from disparate sources. The process began with the careful selection of key variables, including biomarkers, cellular changes, clinical outcomes, and exposure levels, which formed the foundation for data integration. This was followed by data mapping, where information from various studies was aligned with these variables to ensure uniformity across the database.

Additionally, the case group status was considered, considering factors such as acute or chronic sulfur mustard exposure, as well as treatments like CEES (2-chloroethyl ethyl sulfide). This consideration was crucial for categorizing data and understanding the different impacts based on the duration and type of exposure. To encompass the breadth of research, both low-throughput and high-throughput data were incorporated, providing a comprehensive dataset that supports a wide range of analytical possibilities.

### 4.5 Network Visualization

In the development of Mustard-DB, the network visualization tool, Cytoscape, was incorporated to effectively illustrate the complex relationships within the database [8]. This tool enables users to visually explore the intricate connections between different data points, such as the interactions between genes and proteins, the correlation between clinical symptoms and biochemical markers, and the links between various outcomes. By providing a graphical representation of these relationships, the tool aids in the comprehension of the extensive and multifaceted data, facilitating a deeper understanding of the effects of sulfur mustard exposure.

#### 4.5.1 Data Accessibility

Mustard-DB offers users the flexibility to download the full results in Excel format, providing a familiar and versatile means of data analysis.

## 5 Results

### 5.1 Systematic search

In our comprehensive systematic search of 787 studies, we analyzed research related to eyes, lungs, and skin. Specifically, we screened 257 articles on eyes, 408 on lungs, and 541 on skin. Additionally, we considered 65 articles that covered two or all three of these organs. Beyond this, we categorized studies related to other body organs separately. Our analysis revealed 8 types of diseases across 9 organisms and 29 different tissue types. We extracted 10 statistical tests and 37 laboratory techniques, involving a total of 3754 biological variables. Furthermore, we extracted 175 correlations related to variable types in 4 specific tissue types across 4 different diseases. The PRISMA flowchart depicting the outcomes of our systematic search is presented in Figure 2.

### 5.2 Ontologies as patterns

Sixteen ontologies were specifically created for disease types across twenty-two organisms. Additionally, there were 215 ontologies related to various sample types, three types of studies, and sixteen types of experiments, all of which were considered essential characteristics of the research. These studies incorporated one or more variables from a total of 19,772 ontologies, each officially represented by a symbol falling into one of six categories: transcript, protein, metabolite, cell, tissue, and phenotype. Furthermore, the evaluation process involved 210 techniques, twenty types of statistical tests, and three types of difference markers (increase, decrease, or no change).

### 5.3 Database Management Panel

The database management panel consists of three key sections. First, the main settings allow configuration of database parameters, including “Contact Us” messages, database resets, and graph settings. Second, the ontology definition section loads standard variable names across six predicted categories (gene, protein, metabolite, cell, tissue, and phenotype). It also organizes information from studies into differentials and correlations. Rigorous data entry conditions ensure accuracy. Finally, the third section handles ontology information loading prior to data entry Figure 3.

**Figure 3.**
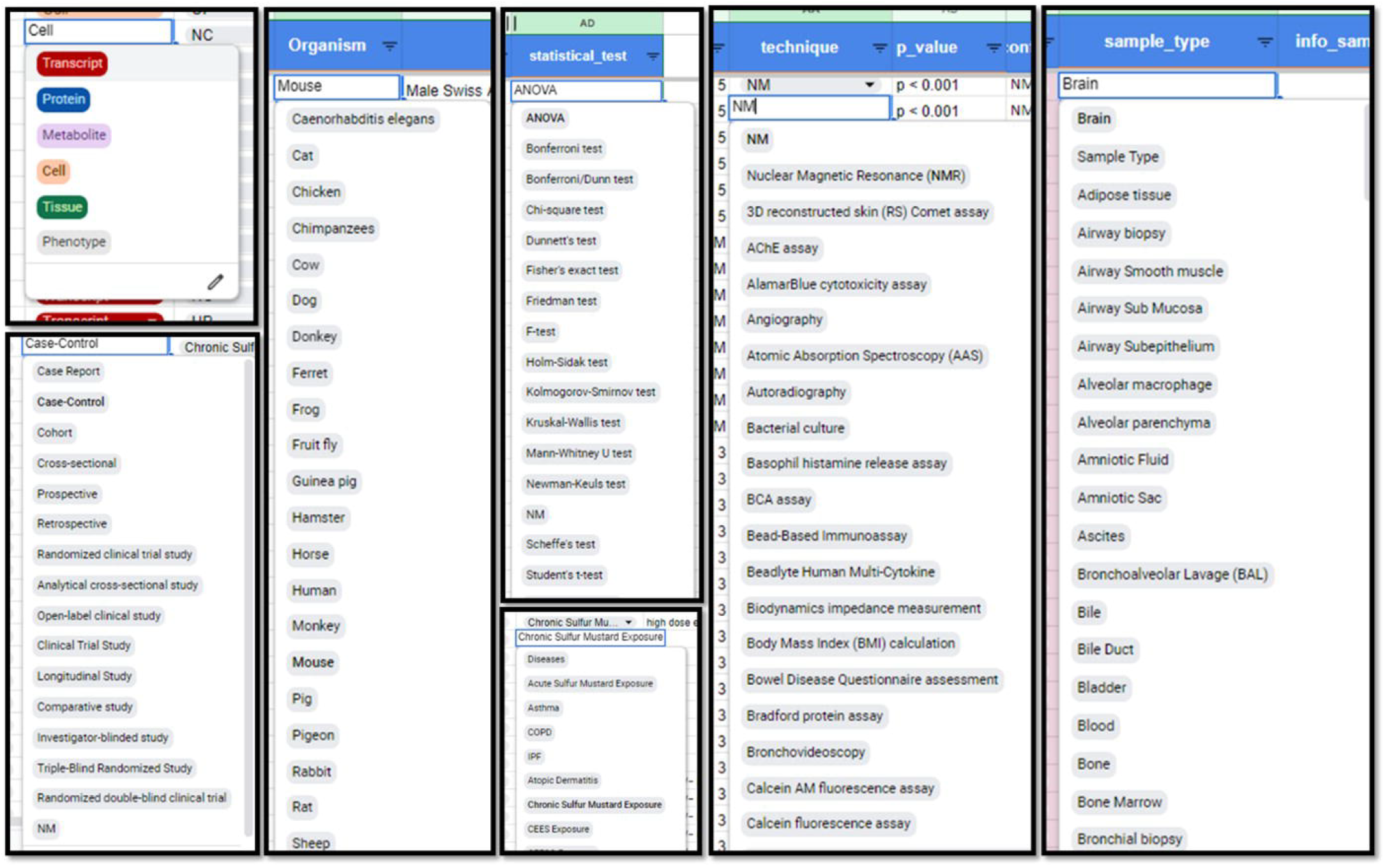
Desing of predefined ontologies in google sheet to increase the uniformity and accuracy of data entry.

**Figure 4.**
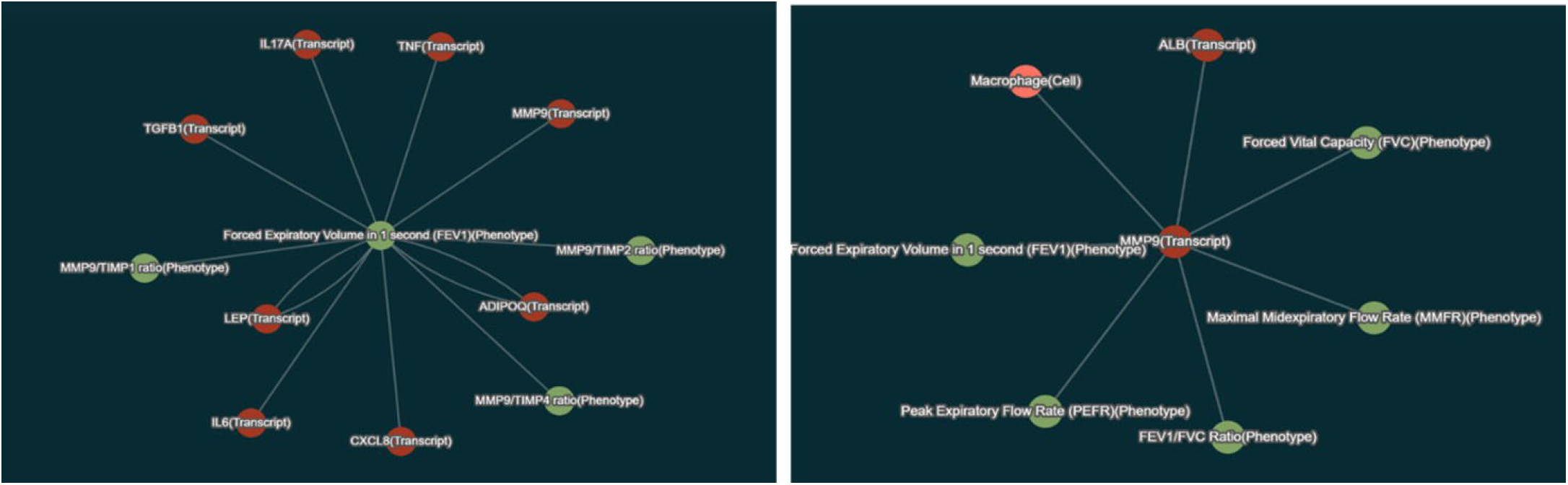
Graphical visualization of relationships between different parameters for FEV1 and MMP9.

### 5.4 Web interface and technical specifications

After providing the necessary infrastructure, the main search page was designed including a simple search box and advanced search bottom so that the users can perform a detailed search by selecting more options from the relevant section. They can investigate queries such as molecules, cells, and phenotypes by entering an intended component name (e.g. “interleukin 1b”), its official symbol (e.g. “IL1B”) or any other alternate name as a query.

By selecting differential or correlation, users can search for queries in the related data. The corresponding table design enables users to view and retrieve the data by searching for desired values in each field. In addition to advanced filtering, data sorting, and outputting possibility, we utilized DataTable to provide easier control and faster performance compared to normal HTML tables (Supplemental File 2).

## 6 Conclusion

The Mustard-DB project represents a significant milestone in consolidating and organizing information from experiments investigating cellular and molecular changes associated with mustard gas exposure. By curating data from diverse sources, this comprehensive database provides valuable insights into the relationship between these changes and cellular, molecular and phenotypic outcomes. It was adjusted to use by researchers across various specialties as an online international and up-to-date database.

The Mustard-DB’s approach emphasizes rigorous data verification by multiple ontologies and specific data structure design. Additionally, the software architecture ensures efficient data entry and provides a user-friendly interface. The Mustard-DB database primarily focuses on articles related to eye, lung, skin, and other organs, meticulously screening and validating the information. It serves as a critical platform for handling substantial data extracted from articles, categorized into differentials and correlations.

The Mustard-DB project addresses the challenges posed by large and diverse datasets; by providing a centralized repository of curated articles, it offers researchers a valuable resource for validation and data entry, aiming to facilitate meaningful research and enhance our understanding of mustard gas-related damage. Its dual focus on differentials and correlations ensures a comprehensive view of the cellular, molecular and phenotypic changes associated to mustard gas exposure. As scientific inquiry continues, Mustard-DB stands as a testament to collaborative efforts, bridging knowledge gaps and advancing our understanding of this critical area of study.

## 7 Ethics

The protocol for patient and sample selection in this study was granted ethical approval by the esteemed ethics committee of Baqiyatallah University of Medical Sciences (IR.BMSU.REC.1399.259).

## 8 Author contribution statement

A.A. and F.R. (co-first authors): Study design, ontology design, QC, data extraction, manuscript writing; M.A.: Data extraction, QC; A.C.: Data extraction, QC; H.B.: Study design; A.G.: Study design; M.A.: Software dev. control, study design; S.G.: Software architecture and development; S.A.J.: Study design, data extraction, ontology design, manuscript writing, and supervision; M.G.: Study design, process control.

## 10 Statement

During the preparation of this work the authors used Microsoft Copilot for assistance in text editing. After using this tool, the authors reviewed and edited the content as needed and take full responsibility for the content of the published article.

## 11 Acknowledgments

The authors acknowledge the support of the Chemical Injuries Research Center, Systems Biology and Poisonings Institute, Baqiyatallah University of Medical Sciences.

## References

1. Amini H, Solaymani-Dodaran M, Ghanei M, Abolghasemi J, Salesi M, Vahedian Azimi A, et al., A 39 Year mortality study of survivors exposed to sulfur mustard agent: A survival analysis, Heliyon. 10 (2024) e24535. 10.1016/j.heliyon.2024.e24535

2. Panahi Y, Jadidi-Niaragh F, Jamalkandi SA, Ghanei M, Pedone C, Nikravanfard N, et al., Immunology of Chronic Obstructive Pulmonary Disease and Sulfur Mustard Induced Airway Injuries: Implications for Immunotherapeutic Interventions, Curr Pharm Des. 22 (2016) 2975–96. 10.2174/1381612822666160307150818

3. Ghanei M, Harandi AA, Molecular and cellular mechanism of lung injuries due to exposure to sulfur mustard: a review, Inhal Toxicol. 23 (2011) 363–71. 10.3109/08958378.2011.576278

4. Ghabili K, Agutter PS, Ghanei M, Ansarin K, Panahi Y, Shoja MM, Sulfur mustard toxicity: history, chemistry, pharmacokinetics, and pharmacodynamics, Crit Rev Toxicol. 41 (2011) 384–403. 10.3109/10408444.2010.541224

5. Elsman EBM, Baba A, Offringa M, committee P-Cs, PRISMA-COSMIN 2024: New guidance aimed to enhance the reporting quality of systematic reviews of outcome measurement instruments, Int J Nurs Stud. 160 (2024) 104880. 10.1016/j.ijnurstu.2024.104880

6. Pang Z, Lu Y, Zhou G, Hui F, Xu L, Viau C, et al., MetaboAnalyst 6.0: towards a unified platform for metabolomics data processing, analysis and interpretation, Nucleic Acids Res. 52 (2024) W398–W406. 10.1093/nar/gkae253

7. Zheng J, Harris MR, Masci AM, Lin Y, Hero A, Smith B, et al., The Ontology of Biological and Clinical Statistics (OBCS) for standardized and reproducible statistical analysis, J Biomed Semantics. 7 (2016) 53. 10.1186/s13326-016-0100-2

8. Franz M, Lopes CT, Fong D, Kucera M, Cheung M, Siper MC, et al., Cytoscape.js 2023 update: a graph theory library for visualization and analysis, Bioinformatics. 39 (2023) 10.1093/bioinformatics/btad031

