## Supplemental File 2 for "Mustard-DB: The Database of Cellular, Molecular, and Phenotypic Data in Sulphur Mustard Exposure"

This supplementary file provides an overview of the data currently hosted on our website, which serves as a foundational resource for our research. As of the latest update, the website contains a total of 1167 data entries. These entries are categorized into two main types: correlation data, which comprises 129 entries, and differential data, which makes up the remaining 1038 entries. This dataset is part of an ongoing effort to build a comprehensive database, with continuous uploads being made to enhance its scope and utility for future analyses. The data presented here offers a snapshot of our current holdings and highlights the potential for further exploration and research.

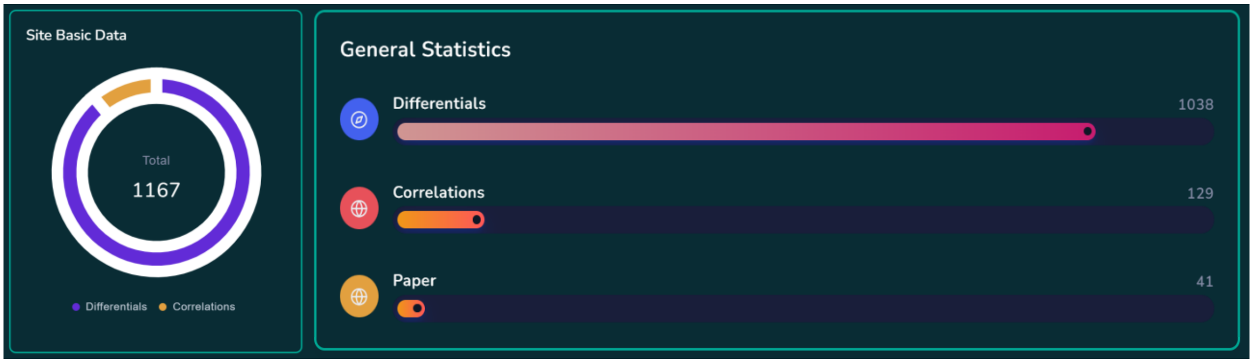


The database encompasses a variety of sample types, including bronchoalveolar lavage (BAL), blood, corneal tissue, liver, lung biopsy, and several others, as illustrated in the accompanying figure. The distribution of correlation and differential data across these sample types is also visualized.

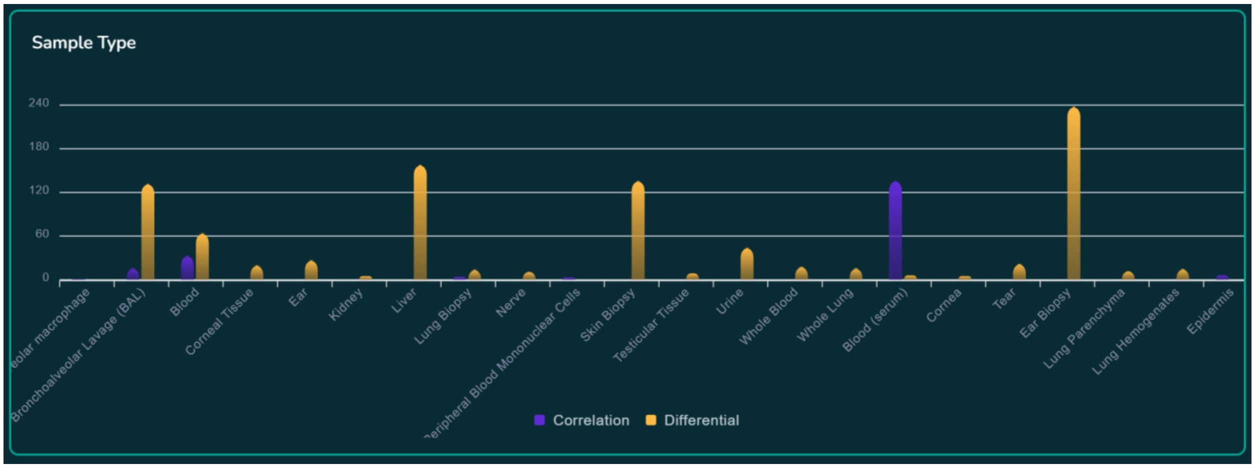


The data table illustrates the number of data points available for each organism, categorized by various biological sample types. Humans, mice, and rats are the most represented organisms, with data available across multiple sample types.

| Organism | Bronchoalve... | Blood | Corneal Tis... | Ear | Kidney | Liver | Lung Biopsy | Nerve | Skin Biopsy | Testicular... | Urine |
|---|---|---|---|---|---|---|---|---|---|---|---|
| Chicken | 0 | 0 | 0 | 0 | 0 | 0 | 0 | 8 | 0 | 0 | 0 |
| Guinea pig | 0 | 0 | 0 | 0 | 0 | 3 | 0 | 0 | 0 | 0 | 0 |
| Human | 48 | 23 | 20 | 0 | 0 | 0 | 0 | 0 | 0 | 9 | 0 |
| Mouse | 30 | 29 | 0 | 27 | 0 | 118 | 0 | 0 | 68 | 0 | 44 |
| Pig | 0 | 0 | 0 | 0 | 0 | 0 | 0 | 0 | 68 | 0 | 0 |
| Rabbit | 0 | 0 | 0 | 0 | 0 | 0 | 0 | 0 | 0 | 0 | 0 |
| Rat | 54 | 12 | 0 | 0 | 5 | 37 | 14 | 3 | 0 | 0 | 0 |

The tables presented here are part of the comprehensive data collection available on our Mustard database. They provide insights into the distribution of data across various biological samples and diseases. For a deeper dive into these datasets and additional analyses, please visit the "Statistics" page on our website. This resource offers detailed information and further breakdowns of the data, allowing users to explore the full scope of our database and its potential applications.
